# RAPMS 2.0 Improves Specificity and Throughput for Proteomic Identification of RNA Binding Proteins

**DOI:** 10.1101/2025.11.23.690051

**Authors:** D. Honson, W. Deng, M. Blanco, J. Sha, K. Plath, J.A. Wohlschlegl, M. Guttman, D. Majumdar

## Abstract

RNA-protein complexes are critical factors in development, homeostasis, and disease. RNA proteomics methods are essential for characterizing these complexes but suffer from high levels of background, which hinders identification of ribonucleoprotein (RNP) components. Here, we present RNA Antisense Purification followed by Mass Spectrometry 2.0 (RAP-MS 2.0), an updated version our original RAP-MS protocol with innovations in bead preparation, RNA capture, and peptide purification. RAP-MS 2.0 has lower background than our original protocol, and allows lysate to be reused to capture multiple RNAs. We demonstrate that RAP-MS 2.0 recapitulates known RNPs for 7SL, 7SK, RMRP, U1, U2, U6, U7, and Xist. Additionally, we use RAP-MS 2.0 to identify novel RNA-protein interactions between Xist and TREX components and U1 with FET family transcriptional regulators.

## INTRODUCTION

RNA-protein complexes are ubiquitous regulators of RNA biogenesis and gene expression. Even before transcription begins, the ribonucleoprotein (RNP) 7SK queues RNA polymerase II for transcription by regulating phosphorylation.^1^ During transcription, RNA binding proteins (RBPs) in the spliceosome, the exon-junction complex, and cleavage and polyadenylation complexes help process and stabilize the nascent transcript.^2^ RBPs escort mRNAs out of the nucleus and assemble them with the ribosome, the largest and most abundant RNP in the cell.^2^ For non-coding transcripts, such as small nuclear RNAs (snRNAs), RBPs escort the RNA to cellular compartments that assemble the mature RNP.^3^ The centrality of RNA-protein complexes to these processes underscores the importance of biological methods to characterize RNA-protein interactions.

RNA proteomics methods were developed to enumerate the components of RNA-protein complexes. The family of methods employs liquid chromatography mass spectrometry (LCMS) to identify protein binding partners of specific RNAs. The proteins can be isolated by purifying the target RNP, selectively degrading the target RNA to release bound proteins, or employing proximity biotinylation in living cells.^4,5^ These approaches have elucidated the protein components of numerous RNPs that were inaccessible to older methods.^6–13^ A limitation of RNA proteomics methods, however, is that they purify large amounts of non-specific background along with RBPs that genuinely bind the target. As such, the datasets require extensive curation to remove false positives.

Specificity in RBP proteomics is particularly challenging in the characterization of dynamic, accessory, or regulatory RBPs that function outside of stable, mature RNPs. Core RNP components bind stably, crosslink efficiently to their RNA partners, and as such are robustly detected by RNA proteomics methods. Many proteins in these datasets, however, are detected at much lower levels, and current methods cannot distinguish non-specific background from bona fide RBPs in the lower abundance population. Nonetheless, these more elusive RBPs are biologically significant.^14–16^ Proteins involved in RNA biogenesis, coordination of RNPs in larger molecular machines, and regulation of RNP activity are among these important factors.

Here, we present RAP-MS 2.0, an adaptation of our original RAP-MS method optimized to reduce background and improve identification of non-core RBPs. Implementation of new approaches to RNA capture involve switching from biotinylated antisense probes to probes covalently linked to a paramagnetic bead, incorporating a capture-elute-recapture strategy to increase specificity, and purifying peptides with carboxylate beads to improve yield (Figure 1a-b). The methodological improvements behind RAP-MS 2.0 yield increased specificity than its antecedent and allow multiple RNA species to be purified from a single lysate. We demonstrate that these modifications improve unambiguous identification of both validated and novel RNA-protein interactions.

**Figure 1.**
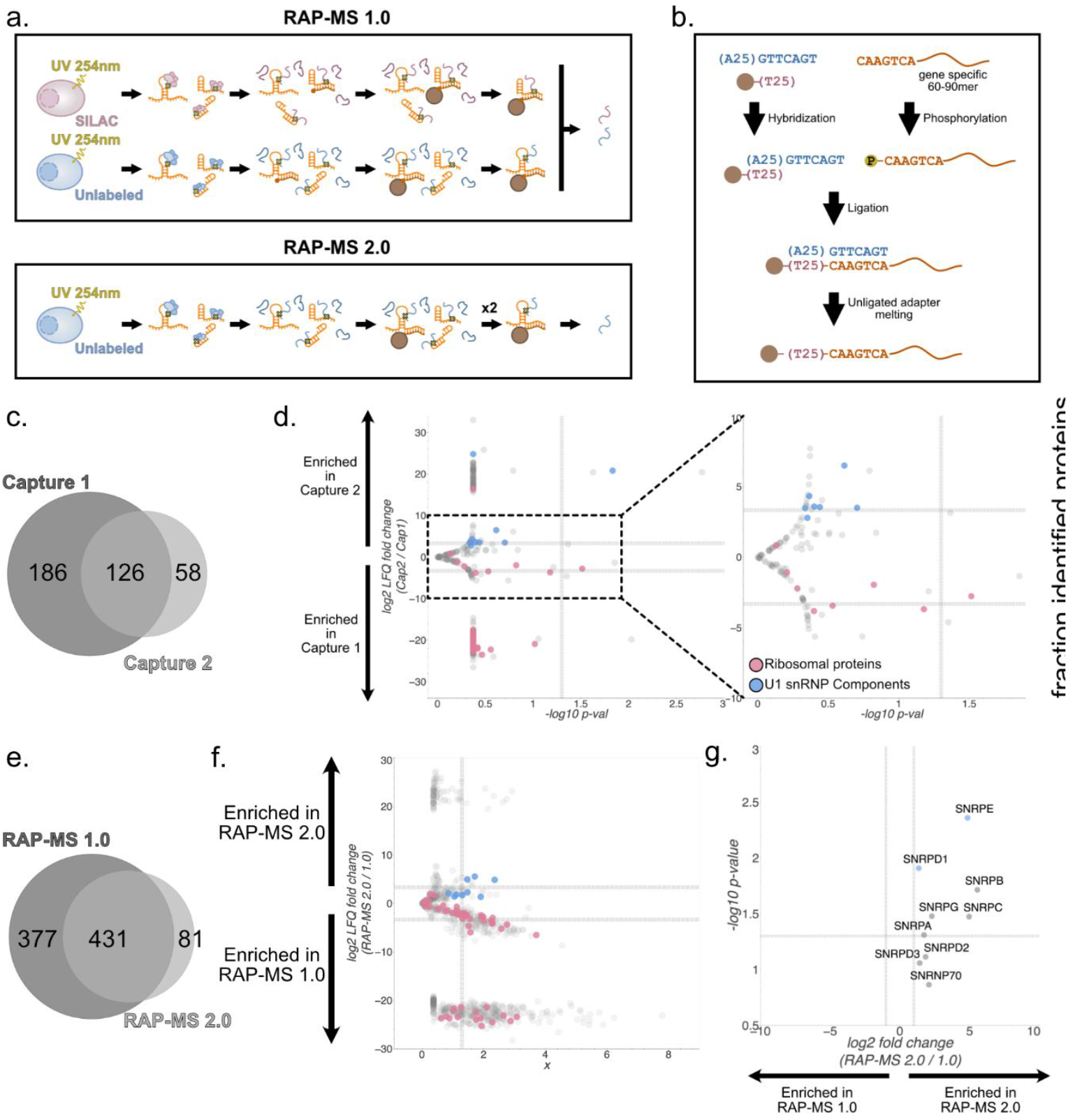
RAP-MS 2.0 uses multiple captures to improve specificity. a, Schematics of RAP-MS 1.0 and 2.0. b, Schematic of probe-bead preparation. c, Unique and shared proteins identified in capture 1 and 2 of U1 snRNA RAP-MS 2.0. d, Volcano plot of proteins enriched in capture 1 and 2 of U1 snRNA RAP-MS 2.0. Horizontal dotted lines represent 2-fold or −2-fold changes in LFQ intensity. Ribosomal proteins (background) are indicated in pink, and U1 snRNP components (signal) are in blue. The right panel is an inset of the left focusing only on proteins identified in both captures. e, Unique and shared proteins identified in RAP-MS 1.0 and 2.0 for U1 snRNA. f, Volcano plot of proteins enriched in RAP-MS 1.0 and 2.0 of U1 snRNA. Vertical dotted lines represent 2-fold and −2-fold changes in LFQ intensity, and the horizontal dotted line represents Students T-test p = 0.05. Ribosomal proteins are indicated in pink, and U1 core snRNP components in blue. g, Inset of panel f showing only U1 snRNP components. Seven of the ten components that differed significantly (p < 0.05) by more than 2-fold are indicated in blue. All that significantly differed were enriched in RAP-MS 2.0.

## RESULTS

### Comparison between RAP-MS and RAP-MS 2.0

RAP-MS 2.0 is based on the original RAP-MS protocol (RAP-MS 1.0, from here forward) but has several innovations. In RAP-MS 1.0, two cell cultures are grown: one metabolically labeled with heavy amino acids (SILAC) and another in standard media (Figure 1a).^17,18^ The RNA of interest is purified from the SILAC culture, and a well-characterized control RNA (such as the U1 snRNA or 18S rRNA) is purified from the unlabeled cells. Both cultures are crosslinked by UV irradiation and lysed in a protein denaturing buffer. Biotinylated DNA probes against the target RNAs are added to each lysate and recovered with streptavidin beads. RNA-bound proteins are eluted with a nuclease, then desalted for LCMS through TCA precipitation.

RAP-MS 2.0 updates four main aspects of the RAP-MS 1.0 protocol (Figure 1a). First, biotinylated probes and streptavidin beads are replaced with a novel bead preparation in which antisense probes are covalently linked to paramagnetic oligo-dT beads (Figure 1b). Second, the new beads allow for the targeted RNA to be non-destructively eluted with heat and recaptured, thereby improving specificity of the RNA purification. Third, the TCA precipitation has been replaced with a “Single-Pot, Solid-Phase-Enhanced Sample Preparation (SP3) to improve protein yields from very low concentration samples.^17^ Finally, the improvements in specificity conferred by the new bead preparation and multiple capture strategy have allowed us to remove SILAC labeling from the protocol.

RAP-MS 2.0 beads are prepared through a sticky-end ligation strategy (Figure 1b). In short, a DNA splint hybridizes to the oligo-d(T)25 sequence with an overhang sequence. The overhang corresponds to a common sequence on the 5’-end of all antisense probes. Hybridization of the antisense probes with the overhang of the splint allows ligation between the antisense probes and the oligo-d(T)25 sequence that is covalently linked to the beads. After establishing the bead preparation protocol, we assessed whether eluting and recapturing the RNA would improve specificity.

We performed RAP-MS 2.0 on U1 snRNA, the RNA component of the U1 snRNP complex, chosen for its abundance and well characterized proteome.^19–23^ Mouse embryonic stem cell (mESC) lysate was exposed to RAP-MS 2.0 beads carrying probes against U1. Captured RNA was eluted with heat and 10% of the eluent was reserved. The remaining 90% was exposed to fresh beads and the elution was repeated. Protein was purified by each capture and analyzed by LCMS. Comparing capture 1 and capture 2, the total number of proteins decreased between the captures, as might be expected from any repeated enrichment, but, importantly, several proteins were exclusively identified in each capture: 186 proteins were exclusively detected in capture 1, compared to 58 exclusive to capture 2 (Figure 1C). This suggests that the high complexity of capture 1 masked low abundance proteins, as conventional Data Dependent Acquisition (DDA) is biased to highly abundant ions. The proteins specific to capture 2 were likely present in the capture 1 sample but were too poorly represented in capture 1 to be robustly detected by DDA. While the second capture might have resulted in some loss in yield, the increased enrichment provided a greater opportunity to identify and reproducibly quantify these less abundant signals.

As U1 forms a well-characterized complex with spliceosomal proteins, we first quantitated the relative capture efficiency of these proteins between the first and second capture (Fig. 1d). In total, we identified eight of the ten core U1 snRNP members (U1-70K, U1-A, U1-C, Snrpb, Snrpd1, Snrpd2, Snrpd3, and Snrpg).^24–26^ Of those, Snrpd1 was exclusively identified in capture 2, indicating that the second capture improved coverage of the U1 snRNP. Six of the seven U1 snRNP factors found in both captures 1 and 2 were enriched by greater than ten-fold in capture 2. Taken together, in the context of the U1 snRNP, adding the stringency of a second capture round improved both sensitivity and specificity of the measurement.

To evaluate specificity, we analyzed the enrichment of ribosomal proteins in each capture (Figure 1d, S1a). Background in RAP-MS experiments is defined as proteins that are not covalently bound to the target RNA but that nonetheless appear in the final eluent. As U1 is not translated, we consider any ribosomal proteins in the sample to be non-specific noise. Overall, 29 of 38 detected ribosomal proteins identified were exclusively found in capture 1 (Figure S1A). Only one was specific to capture 2, and it was only found in one replicate. 8 of 38 ribosomal proteins were identified in both captures, and all but one were enriched in capture 1. The enrichment for U1 core proteins and depletion of ribosomal contaminants in capture 2 confirms that the two-capture strategy improves signal-to-noise ratio in RAP-MS 2.0.

To compare RAP-MS 2.0 with its precursor, RAP-MS 1.0, we captured U1 snRNA using both methods and applied the same analyses as we did with the first and second captures. 377 proteins were exclusive to RAP-MS 1.0, compared with only 81 specific to RAP-MS 2.0 (Figure 1E), in line with the yield differences observed when captures were compared. We examined ribosomal proteins to determine whether the higher signal in RAP-MS 1.0 was due to background or superior identification of RNA binding proteins.

55 ribosomal background proteins were identified between both methods (Figure 1F, S1B). 49 of the 55 ribosomal proteins were enriched in RAP-MS 1.0, of which 16 were exclusively found in RAP-MS 1.0 samples. No ribosomal proteins were exclusive to RAP-MS 2.0, and those that were enriched in RAP-MS 2.0 did not differ significantly. This indicates that the RAP-MS 2.0 method substantially decreases the number of background proteins compared to our original protocol. Notably, the intensities of proteins found by both methods (48.5% of all proteins) correlated well (R^2^ = 0.48, compared to R^2^ = −0.08 when all proteins were included). The high degree of correlation suggested that the methods may agree well on bona fide U1 binding proteins but diverge on background.

To assess the agreement of RAP-MS 1.0 and 2.0 on U1 binders, we examined core components of the U1 snRNP (Figure 1F-G). Both methods identified all ten members of the U1 snRNP. All were enriched in RAP-MS 2.0 relative to 1.0, but only two differed significantly (Student’s T-test p < 0.05, Benjamini-Hochberg FDR = 0.05, Figure 1G). We therefore observe enrichment of well-characterized U1 binders and marked depletion of contaminants, suggesting that RAP-MS 2.0 successfully improves specificity compared to our original protocol and is a method with increased suitability for identifying *bona fide* RNA-protein interactions.

### RAP-MS 2.0 multiplexing by serial captures

A key distinction between RAP-MS 1.0 and 2.0 is that the newer method does not degrade the purified RNA during elution. This distinction allowed for the possibility of significant increases in experimental scale if multiple rounds of RBP identification from could be achieved from the same lysate. RAP-MS experiments can require upwards of 10^8^ cells to obtain high enough levels of crosslinked RNA for proteomic analysis, depending on RNA of interest. The RAP capture, however, is intended to remove a negligible fraction of the total RNA from the lysate. We speculated that flowthrough from the first capture would be nearly as rich in RNA-protein complexes as the original lysate, permitting us to speculate that multiple rounds of capture might be possible.

We therefore tested the reusability of the lysate flowthrough through successive experiments, possible because covalently bound probes are completely removed after the first capture (Figure 2A). Broadening our studies to multiple classes and lengths of RNA, we attempted to capture RBPs bound to a diverse set of non-coding RNAs (XIST, RMRP, 7SK, and 7SL) and spliceosome-associated snRNAs (U1, U2, U6, U7). To do this, we prepared lysate from ~75M mESCs containing Xist under a dox-inducible promoter. We incubated the lysate with RAP-MS 2.0 beads carrying probes against Xist, then magnetically removed the beads. We then incubated the flowthrough from the Xist capture with 7SL-targeting beads. We repeated the process with beads targeting 7SK and U1, capturing a total of four RNAs from a single lysate. Once all first captures were collected, each was processed independently for washes, second capture, elution, and LC-MS analysis. We performed the same workflow on a different lysate to capture U7, RMRP, U6, and U2 (in that order). Each experiment was performed in duplicate, leading to a total of 16 RAP-MS datasets.

**Figure 2.**
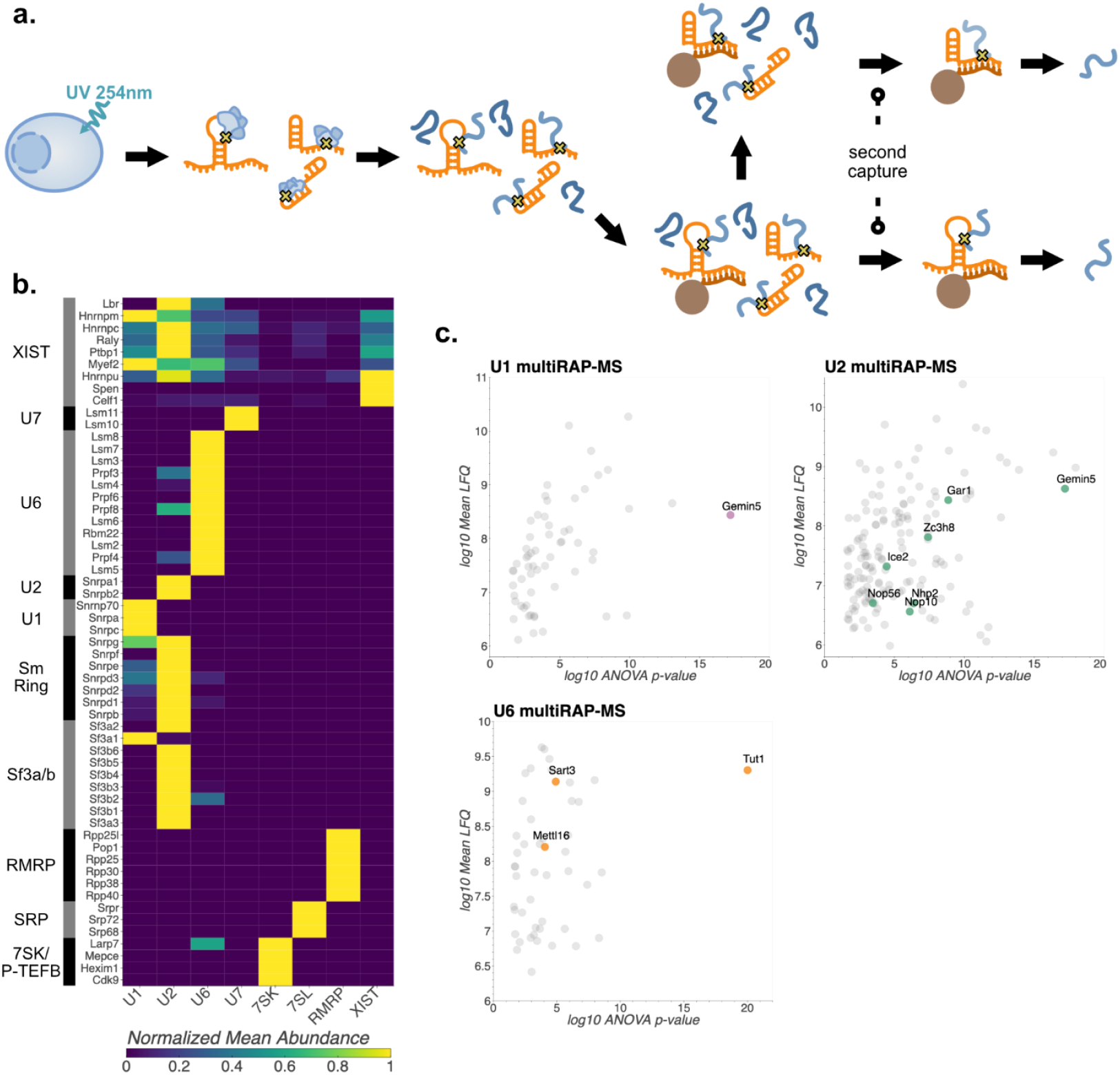
Mulitplexed RAP-MS 2.0 recapitulates known RNP. a, Schematic of multiplexed RAP-MS 2.0 (multi-RAP-MS 2.0). b, Heatmap of core RNP components. Individual proteins grouped by RNP are indicated on the y-axis while multi-RAP-MS 2.0 target is on the x-axis. Color indicates the LFQ intensity values normalized per protein. c, U1, U2, and U6 snRNA multi-RAP-MS 2.0 highlighting biogenesis factors.

The strategy of multiplexed RAP-MS 2.0 (‘multi-RAP-MS 2.0’) performed on successive flowthrough was validated by evaluation of each RNA’s well-characterized proteome. RBPs well known to interact with each RNA identified RBPs members of RNPs for all RNAs targeted (Figure 2B).^18,27–34^ We did not observe any obvious effect of order of capture, and, consistently, RBPs of earlier captures were not observed in later ones (e.g., 7SK-bound proteins were not observed in the U1 dataset). Indeed, U1 and U2, RNAs captured in the final of the four successive captures, both yielded abundant RBP despite being the last captured lysate.

In addition to the expected complexes, examination of RBPs that comprise RNPs associated with several of the captured RNAs revealed some reasonable but unexpected interactions. The SF3a and b complexes are typically considered part of the U2 snRNP.^35,36^ RAP-MS confirmed that most SF3 components were specific to U2, but also indicated interactions between U1 and SF3a1, and U6 and SF3b2 (Figure 2B). This is consistent with previous research showing that SF3a1 binds to U1 stem-loop 4 in pre-spliceosomal complexes and SF3b2 stabilizes U6 on pre-mRNA.^37,38^ These results demonstrate that Multi-RAP-MS 2.0 allows for multiple RNA species to be recovered from a single lysate without compromising the quality of the individual datasets.

Multi-RAP-MS-2.0 for U1, U2, and U6 revealed factors involved in numerous steps of snRNP biogenesis (Figure 2c). Both U1 and U2 showed binding to Gemin5, a component of the SMN complex essential for Sm ring assembly on immature snRNAs.^39^ Additionally, U2 capture revealed three Cajal body proteins (Cofilin, Ice2, and Zc3h8) and five H/ACA snoRNA components (Dck1, Gar1, Nhp2, Nop10, and Nop56). Cajal bodies are the site of snRNA pseudouridylation and scaRNAs, a subclass of snoRNAs found in Cajal bodies, catalyze this post-transcriptional modification. U6 capture recovered two proteins associated with U6 biogenesis: Tut1 which uridylates the 3’-end of immature U6, and Mettl16, which methylates U6 adenosine residues.^40–43^ U6 RAP-MS additionally revealed binding to Sart3. After U6 is ejected from the post-spliceosomal complex, Sart3 binds to the snRNA and facilitates regeneration of the U4/6 di-snRNP.^44^ These results demonstrate that RAP-MS identifies RBPs outside core RNPs and can provide insights into the life cycles of functional RNAs.

A challenge of RNA-proteomics is distinguishing between direct and indirect RNA-protein interactions. This problem is especially acute in cases like the spliceosome, which undergoes multiple conformational and compositional rearrangements with as many as five snRNPs in a single assembly (U1, U2, U4, U5, and U6 in the early pre-catalytic B complex).^45^ To examine whether RAP-MS could correctly assign RNA-protein interactions known from cryo-EM structures, we examined three non-snRNP spliceosomal components that act in the transition from the pre-catalytic (B) to catalytic (B*) spliceosome: Snw1 (Skip), Ppie (CycE) and Syf2.^46^ This transition requires at least three intermediate structures: the early, mature, and late active spliceosomes (B^act^). The precise orchestration of this process is required to position the 2’-hydroxyl group of the branchpoint adenosine for nucleophilic attack on the 3’-hydroxyl of the splicing donor, leading to formation of the lariat.^47^

Snw1 is a splicing factor that enters the spliceosome during the transition from the pre-catalytic B complex to the early B^act^ complex.^46^ Existing structures of the B^act^ complexes have substantial missing densities for Snw1, but it interacts specifically with U2 in the C* spliceosome, the structure that catalyzes the second transesterification reaction.^46^ Using RAP-MS 2.0, we find that Snw1 binds exclusively to U2, as compared to U1 and U6 (Figure 3a-b).

**Figure 3.**
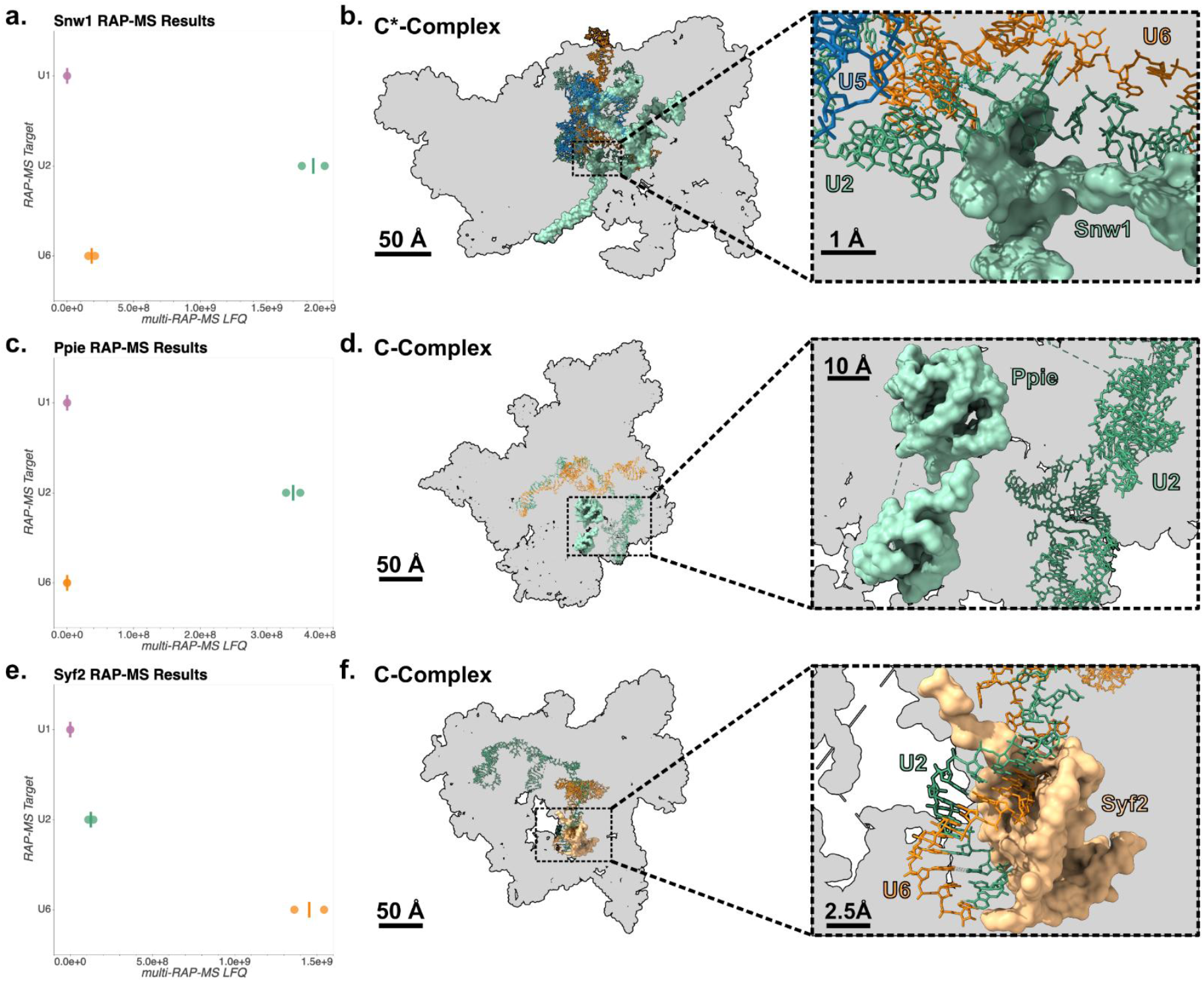
Multi-RAP-MS 2.0 concurs with structural data. a, Multi-RAP-MS intensity of Snw1 for U1, U2, and U6. Snw1 was strongly enriched in the U2 RAP. b, Cryo-EM structure of the C*-spliceosomal complex highlighting the U2-Snw1 interaction (PDB ID 5XJC). c, Multi-RAP-MS intensity of Ppie for U1, U2, and U6. Ppie was strongly enriched in the U2 RAP. d, Cryo-EM structure of the C-spliceosomal complex highlighting the U2-Ppie interaction (PDB ID 5YZG). e, Multi-RAP-MS intensity of Syf2 for U1, U2, and U6. Syf2 was strongly enriched in the U6 RAP. f, Cryo-EM structure of the C*-spliceosomal complex highlighting the U6-Syf2 interaction (PDB ID 5YZG).

Ppie is a peptidylprolyl isomerase also known as cyclophilin E enters the spliceosome in the mature B^act^ complex.^46^ It remains in the spliceosome during the first transesterification reaction in the B* complex and becomes closely associated with the U2 snRNP in the resulting C complex.^48,49^ Consistent with this interaction, RAP-MS exclusively showed an interaction between U2 and Ppie (Figure 3c-d).

Syf2 is a member of the NineTeen Complex (NTC), which enters the spliceosome in the B-complex and guides spliceosomal conformational changes until the completion of splicing.^50^ In existing structures, Syf2 binds closely to the U2/U6 duplex in the C complex, but both RAP-MS showed a specific enrichment of Syf2 over U6 (Figure 3e-f).^49^ This may indicate that Syf2’s association with U6 is more stable than with U2, which could be functionally significant for stabilizing spliceosomal structures. Taken together, direct interactions from dynamic assemblies such as the spliceosome are accurately captured with Multi-RAP-MS 2.0, as measurements of three spliceosomal snRNAs recapitulate the differential interactions of each subcomplex.

### Multi-RAP-MS 2.0 reveals novel ncRNA-protein interactions

Multiplexed RAPMS2.0 also revealed new and previously uncharacterized interactions, six of which we pursued for further validation.: Chtop and Alyref binding to Xist, Nepro binding to RMRP, and Fus, Ewsr1, and Taf15 (FET) binding to U1. These proteins were selected because their RAP-MS signals were comparable to those of core RNP components, and because their annotated functions suggested they might augment the activities of the RNAs they bind.

Many lncRNAs, including Xist, share molecular features with mRNAs such as introns, m7G-caps, and polyA tails.^51^ Unlike mRNAs, however, not all lncRNAs are destined for the cytoplasm. An open question in lncRNA biology is how the cell determines which lncRNAs remain in the nucleus and which are exported. Xist is a classic example of a nuclear lncRNA; it remains bound to the inactive X-chromosome for its entire lifecycle. Unexpectedly, multi-RAP-MS 2.0 revealed that Xist strongly and specifically interacts with two members of the transcription export (TREX) complex: Alyref and Chtop (Figure 4A).^52^ To confirm this interaction, we analyzed iCLIP data for Alyref and Chtop from a previous study, (Figure 4B).^53^ As expected, both proteins bound robustly to Xist compared to a negative control. TREX has functions beyond nuclear export such as influencing splice-site and polyadenylation site choice.

**Figure 4.**
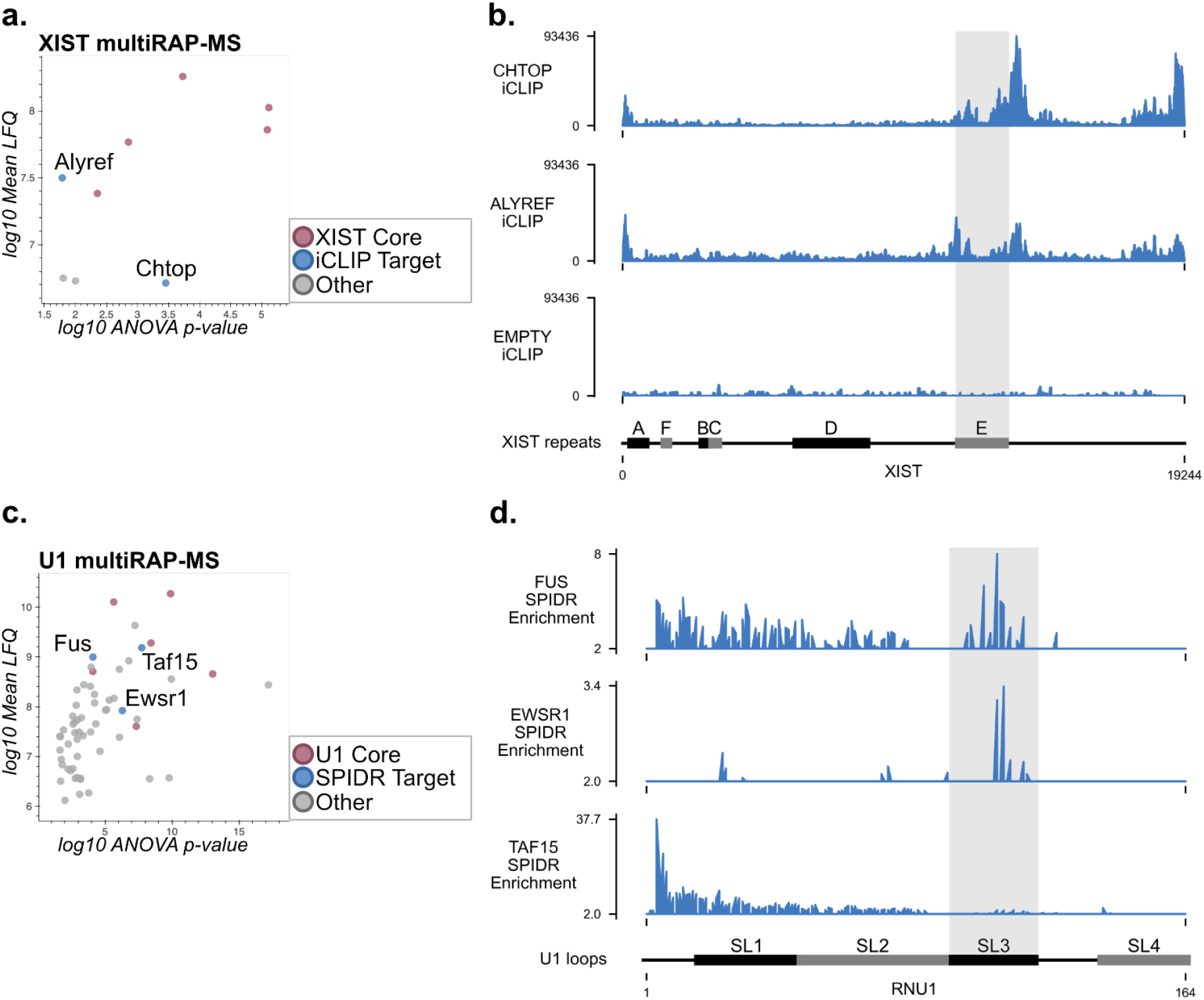
Multi-RAP-MS 2.0 reveals novel RNA-protein interactions. a, Intensity vs ANOVA p-value for significantly differing Xist-bound proteins. In addition to known core proteins, multi-RAP-MS 2.0 reveals TREX components Alyref and Chtop. b, anti-FLAG iCLIP data for Chtop-FLAG, Alyref-FLAG, and Empty-FLAG from Viphakone et al, Mol Cell 2019 over human XIST.^53^ Plotted data are from a bedgraph with 1nt bins. Both TREX components show 5’-, 3’- and E repeat enrichment. c, Intensity vs ANOVA p-value for significantly differing RMRP-bound proteins. In addition to known core proteins, multi-RAP-MS 2.0 reveals the nucleolar protein Nepro. d, CLAP data for Nepro over mouse RMRP and 45S-ITS1. Plotted data are from a bedgraph with 1nt bins. e, Intensity vs ANOVA p-value for significantly differing U1 snRNA-bound proteins. In addition to known core proteins, multi-RAP-MS 2.0 reveals the FET family of transcriptional regulators: Fus, Ewsr1, and Taf15. f, SPIDR enrichment of Fus, Taf15, and Ewsr1 over human U1 snRNA. Data plotted are second read truncations indicating UV crosslinking sites. Fus and Ewsr1 show enrichment over stem-loop 3 (SL3) and Taf15 shows enrichment over the 5’-splice site recognition motif.

Our next targets were Fus, Ewsr1, and Taf15 over the U1 snRNA. Fus, Ewsr1, and Taf15 comprise the FET family of RNA-binding transcriptional activators (Figure 4C).^61^ Loss of U1 and FET proteins share many phenotypes: reduced transcriptional activation, changes to alternative splicing, and premature cleavage and polyadenyation.^61–63^ Consistent with these shared functions, previous research has suggested that the U1 snRNP interacts with both Fus and Taf15.^64–68^ To our knowledge, U1’s interaction with Ewsr1 has not been previously reported.

To map FET protein binding sites to U1, we reanalyzed our previously published data from Split-Pool Identification of RBP Targets (SPIDR) (Figure 4D).^69^ In SPIDR, magnetic beads carrying both an antibody and a unique oligonucleotide sequence (an antibody ID) are exposed to UV-crosslinked lysate. After immunoprecipitation, captured RNA is reverse transcribed. The cDNA and antibody ID are labeled by split-pool ligation of oligonucleotide tags. After sequencing, individual beads are identified by their split-pool barcodes, and cDNA is assigned to antibodies using their antibody IDs. SPIDR revealed binding of Taf15 near the 5’-splice site recognition motif, and of Fus and Ewsr1 to stem-loop 3. Notably, the Fus binding-site matches the results of a recent paper that employed CLIP and NMR to solve a partial Fus-U1 structure.^66^ Recent research has also indicated that the interaction between Fus and U1 may be relevant in the development of Amyloid Lateral Sclerosis (ALS).^66,70^

## DISCUSSION

The development of proteomics approaches to identify RBPs led to a tremendous expansion of the number of putative RBPs in the scientific literature.^71–77^ While the protein components of RNA-protein complexes are central to many enduring biological questions, methodologies to discover RBPs are often noisy, an issue common to many RNA technologies.^78^ In this study, we present RAP-MS 2.0, a significant enhancement of our original methodology of RBP capture by RNA pulldown. The key innovations of the protocol are 1) using probes covalently linked to magnetic beads in place of a biotin-streptavidin linkage, 2) employing heat rather than nuclease elution to enable a dual-capture strategy and multiplexing, 3) using label free quantification rather than SILAC for LC-MS analysis, and 4) using carboxylate beads rather than TCA precipitation for protein and peptide recovery.

Our novel bead preparation allowed us to elute the target RNA with heat while keeping the probe bound to the beads. This enabled a dual capture strategy that increased stringency and specificity. We found the sequential capture strategy improves stringency by reducing recovery of off-target RNAs and proteins. This generalizable approach has been shown to improve purity of yield in other contexts.^79,80^ We found that the added stringency of this step clearly removes ribosomal contamination from the spliceosomal U1 snRNP, as compared to a single capture with metabolic labeling. The background reduction in RAP-MS 2.0 has allowed us to confidently identify ‘accessory’ components of RNA-protein complexes. We demonstrated this advantage by identifying snRNA-binding proteins that represent the whole of the snRNA lifecycle, from their biogenesis to the specific snRNA-protein binding dynamics in the spliceosome, and finally through their recycling after release from the spliceosome.

The presence of the free dT stretch on RAP-MS 2.0 beads and the larger amount of probe could have led to more background than the biotin-streptavidin strategy by capturing all poly-adenylated RNAs. Were this the case, we would expect to see high levels of ribosomal proteins due to mRNA contamination. This was not the case. Ribosomal proteins were rarely enriched in any of our captures, and in fact our updated protocol shows substantially less ribosomal contamination than the original biotin-streptavidin strategy.

The ability to completely remove beads and probes from the lysate also allowed multiplexing by capturing new RNAs from the flow-through of the first capture. This advancement, along with the use of label-free quantification (over SILAC), greatly improved throughput and feasibility of these studies. Using these strategies, we present a new and optimized methodology for the routine discovery of new RBP interactomes for any given RNA. We also demonstrate the power of this method in proof-of-principle experiments on eight well-studied RNAs, revealing several novel interactions.

Using the multiplexing strategy along with RAP-MS 2.0’s improved sensitivity, we identified novel RNA-protein interactions that comprise otherwise well-studied RNA complexes. The binding of TREX components to Xist adds complexity to our understanding of why some RNAs are retained in the nucleus. Although TREX binding is essential for export of mRNAs from the nucleus, it does not accomplish this function when associated with Xist. This may imply that TREX activity is contextual based on the RNA bound, or that the RBPs retaining Xist on the inactive-X overwhelm TREX-mediate export. Additionally, the binding of FET family transcriptional regulators to U1 contributes to long-standing observations about the role of U1 in transcriptional licensing and termination. Although Fus binding to U1 has been previously implicated in U1’s regulation of premature cleavage and polyadenylation, Taf15’s role in U1 biology has not been extensively studied.^64,81^ Moreover, the interaction between Ewsr1 and U1 has not, to our knowledge, been previously reported. Crucially, these interactions were validated by complementary protein-capture with RNA crosslinking and sequencing, Additionally, future application of RAP-MS 2.0 to more poorly understood ncRNAs such as repeats (such as LINEs, SINEs, and satellites), vault RNAs, and lncRNAs may deepen our understanding of the biology of these molecules.

## LIMITATIONS

A consequence of the reduced background of RAP-MS 2.0 may be a small but significant reduction in signal. This may explain the failure to comprehensively identify all RNP components in all replicates. Care should be taken to optimize cell input counts, RNA fragmentation, and probe selection to limit this error. A challenge that remains in RNA-proteomics is the identification of protein partners of very low abundance RNAs, such as nascent mRNAs and many lncRNAs. This limitation is largely due to crosslinking efficiency. UV light covalently links RNAs and proteins at a very low rate, and more efficient crosslinkers such as formaldehyde generally do not exclusively link direct RNA-protein interactions.

RAP-MS 2.0 does not solve this problem, but the high stringency of RAP-MS 2.0 will be required when better crosslinking approaches are developed. Each RNA will carry more protein, meaning that non-specific RNA contamination will lead to even higher levels of off-target proteins in the MS data. RAP-MS 2.0’s two-capture strategy should reduce the contamination problems that higher crosslinking efficiency would introduce.

## METHODS

### Cell culture

#### Mouse embryonic stem cell culture

RAP-MS and CLAP experiments were performed on female mouse embryonic stem cells heterozygous for dox-inducible *Xist* (TX1072; gift from E. Heard lab) as previously described.^82^ Briefly, TX1072 mESCs were grown on gelatin-coated plates in serum-containing ES cell medium (high glucose DMEM (Gibco, Life Technologies), 15% FBS (Omega Scientific), 2mM L-glutamine (Gibco, Life Technologies), 1mM sodium pyruvate (Gibco, Life Technologies), 0.1mM β-mercaptoethanol, 1000 U/ml leukemia inhibitory factor (LIF, Chemicon), and 2i (3 μM Gsk3 inhibitor CT-99021, 1 μM MEK inhibitor PD0325901). Cell culture medium was replaced every 24 hours.

For RAP-MS experiments targeting *Xist*, TX1072 cells were treated with 2µg/mL doxycycline (Sigma) for 72 hours. Media containing doxycycline was replaced every 24 hours.

#### Overexpression of Halo-tagged constructs

For CLAP, N-terminal Halo tagged Nepro was expressed from a plasmid containing a CAG promoter. TX1072 cells were lifted, pelleted, and counted. 400,000 cells for each transfection were transferred to 10mL ES cell medium in gelatinized 10-cm plates. Lipofectamine 2000 was used to transfect the cells immediately after plating. 18µg plasmid DNA in 450µl Opti-MEM was mixed with 36µl Lipofectamine 2000 in 450µl OptiMEM and incubated for 5 minutes at room temperature for each set of duplicates. 450µl of the reaction was added to each 10-cm plate. Cells were collected approximately 24 hours after transfection.

#### UV crosslinking

Cells were removed from the incubator and washed with ice-cold PBS. The PBS was aspirated, the lids of the plates removed, and the plates transferred to a UV crosslinking chamber. For CLAP, cells were treated with 2.5×10^5^ µJ/cm^2^ 265nm UV light. For RAP-MS, cells were treated with 6.0×10^5^ µJ/cm^2^ 265nm UV light. Cells were immediately removed from the crosslinker and ice-cold PBS was added to the plates. Cells were scraped and collected in conical tubes, then spun 3 minutes, 330g, 4°C. The supernatant was removed, cells were resuspended in 1mL PBS per 50M cells, and 1mL cell suspension was transferred to Eppendorf tubes. Cells were spun 3 minutes, 1000g, 4°C. The supernatant was removed, cells were snap-frozen in liquid nitrogen, and stored at −80°C until use.

### RAP-MS 2.0

*Probe design* A Python package for probe design is available through pip, which contains a link to the documentation (https://pypi.org/project/probeutils/). Probes for short RNAs (U1, U2, U6, 7SK, 7SL, and RMRP) were 60nt long (53nt antisense plus 7nt of adapter) and for Xist were 90nt long (83nt antisense plus 7nt adapter). FASTA files for each RNA were downloaded from NCBI nucleotide. All probes had a 5’-CAAGTCA adapter. For Xist, non-specific probes were removed by querying BLAT (maximum of 25 off-target matches in mm39) or Dfam (probes containing any *Mus musculus* annotated repeats). No such filtering was performed for the short RNAs, as many have unannotated pseudogenes in mm39 that are not causes for concern. C**omplete probe sequences are available in supplementary table**. All probes were ordered from Integrated DNA Technologies (IDT) with standard desalting.

#### Bead preparation

Probes for each RNA were pooled in equimolar ratios to 100µM final. For each RAP, 10 nmol probes were phosphorylated with phosphonucleotide kinase (NEB) in 150µl reactions for 30 minutes at room temperature.

500µl oligo d(T)25 beads (NEB) were washed twice in 500µl Oligo dT Wash Buffer (50 mM HEPES pH 7.5, 300 mM NaCl, 2.5 mM EDTA, pH 8.0, and 0.1% Triton-X 100) then resuspended in 250µl Oligo dT Binding Buffer (50 mM HEPES, pH 7.4, 500 mM LiCl, 2.5 mM EDTA, pH 8.0, and 0.1% Triton-X 100). 10 nmol 1mM polyA bottom (TGACTTGA_25_) was added to the beads and the mixture was incubated 30 minutes, 25°C, shaking at 1000rpm. The beads were then washed three times in 500µl Oligo dT Wash Buffer.

The final wash was removed and the beads were resuspended in the 150µl phosphorylated probe reaction plus 150µl 2x Quick Ligation Buffer (2.5x NEB 5x Quick Ligation Buffer, 0.625x NEB 2x Instant Sticky-End Master Mix, and 18.75% 1,2-propanediol (Sigma)). The ligation was incubated 30 minutes, 25°C, shaking at 1250rpm. The beads were then washed three times in 500µl Oligo dT Wash Buffer.

To remove unligated probe and excess adapter, beads were resuspended in 500µl TE Elution Buffer (20 mM Tris-HCl, pH 7.4, 1 mM EDTA, pH 8.0, and 0.1% sodium dodecyl sulfate) and incubated 2 minutes, 95°C, shaking at 1250rpm. This wash was repeated for a total of three TE washes. After the final wash, beads were washed three times in 4M Urea Hybridization Buffer (4M urea, 10 mM Tris-HCl, pH 7.4, 5 mM EDTA, pH 8.0, 500 mM LiCl, 0.5% Triton-X 100, 0.2% sodium dodecyl sulfate, and 0.1% sodium deoxycholate). After the final wash, beads were resuspended in 250µl 4M Urea Hybridization Buffer and left at room temperature until the first capture.

#### Lysate preparation

50M cell pellets were removed from −80°C and thawed on ice for 10 minutes. For U1 RAP-MS experiments comparing captures 1 and 2 and comparing RAP-MS 1.0 and 2.0, 50M cells per replicate were used. For multi-RAP-MS experiments, ~85M cells per replicate were used. Each pellet was resuspended in 1mL RAP-MS Lysis Buffer (10 mM Tris-HCl, pH 7.4, 5 mM EDTA, pH 8.0, 500 mM LiCl, 0.5% Triton-X 100, 0.2% sodium dodecyl sulfate, and 0.1% sodium deoxycholate) and incubated on ice for an additional 10 minutes. Pellets were homogenized by triturating the lysate with an 18 gauge needle, then spinning through Qiashredder columns (Qiagen). Lysates were sonicated with a Branson probe sonicator for 42 cycles, 0.7 seconds on, 2.3 seconds off, 4-5W power. After sonication, lysates were spun 20 minutes, 16000g, 4°C and supernatants were transferred to clean tubes. 1mL of 8M Urea Hybridization Buffer (8M urea, 10 mM Tris-HCl, pH 7.4, 5 mM EDTA, pH 8.0, 500 mM LiCl, 0.5% Triton-X 100, 0.2% sodium dodecyl sulfate, and 0.1% sodium deoxycholate) was added to each milliliter of lysate for 4M urea final. Lysates were then combined and pre-warmed to 42°C for the first capture.

##### First capture

125µl prepared probe-beads were added to the lysate and the mixture was incubated 30 minutes, 42°C, shaking at 1000rpm. The beads were then magnetically separated. For single RAP-MS experiments, the supernatant was discarded. For multi-RAP-MS experiments, the supernatant was transferred to a new tube, magnetically separated for a second time, then transferred to another clean tube. The supernatant was then placed back at 42°C for 5 minutes, the next set of beads was added, and the incubation was repeated.

After removing the supernatant, beads were washed twice in 4M Urea Hybridization Buffer, twice in SDS Wash Buffer (50mM HEPES, pH 7.4, 10% SDS, and 10mM EDTA), and twice in Oligo dT Wash Buffer. Each wash was performed with 500µl buffer for 2 minutes, 37°C, shaking at 1100rpm. The beads were then washed twice in 500µl TE Elution Buffer at room temperature then resuspended in 150µl TE Elution Buffer

To elute RNA, beads in TE Elution Buffer were heated 3 minutes, 95°C, shaking at 1350rpm. The beads were briefly spun to collect condensation then magnetically separated. The supernatant was transferred to a clean tube and the beads were again suspended in 150µl TE Elution Buffer. The heat elution was repeated, 3 minutes, 95°C, shaking at 1350rpm. After brief spinning and magnetic separation, the second eluent was pooled with the first. For capture 1 and 2 comparisons, 30µl of this eluent was reserved as the capture 1 sample.

##### Second capture

The first capture elution was adjusted to 500mM LiCl with 8M LiCl, then 1 volume of 8M Urea Buffer was added. 125µl prepared probe-beads were added to the lysate and the mixture was incubated 30 minutes, 42°C, shaking at 1000rpm. The beads were then magnetically separated. Washes in 4M Urea Hybridization Buffer, SDS Wash Buffer, Oligo dT Wash Buffer, and TE Elution Buffer were performed as in the first capture. RNA was eluted from the beads as described for the first capture.

##### Peptide preparation and LCMS

5µl hydrophobic and 5µl hydrophilic Sera-Mag carboxylate-coated, paramagnetic beads (Cytiva) were washed three times in 500µl LCMS grade water (Fisher) then resuspended in 10µl water. 300µl elutions were reduced and alkylated with 5mM TCEP and 1.2mM IAA 20 minutes, 25°C, shaking at 1200rpm. 10µl washed beads were added to each sample along with 350µl 100% ethanol. The mixture was incubated on a rotator 10 minutes, room temperature to precipitate proteins onto the beads. The beads were then washed three times with 500µl 80% ethanol. After the final wash was removed, the beads were allowed to dry and then were resuspended in 30µl 50mM TEAB. 1µl Lys-C and 2µl Trypsin were added to the beads, and proteins were digested overnight at 37°C.

To recover the peptides, 1mL acetonitrile (ACN) was added to the beads and the suspension was incubated on a rotator 10 minutes, room temperature. The beads were then washed three times in 500µl ACN. After the final wash was removed, the beads were resuspended in 30µl 2% DMSO to elute the peptide. The beads were magnetically separated and the supernatant transferred to a clean tube. Peptides were lyophilized in a speed-vacuum, then resuspended in 10µl 5% formic acid for LCMS injection.

#### LCMS was performed on an Orbitrap Eclipse and peptides were analyzed using MaxQuant

##### Data analysis

In all analyses, known contaminants such as keratins and trypsin that were flagged by MaxQuant were removed. For the capture 1/capture 2 and RAP-MS 1.0/2.0 analyses (Figure 1c-g), label-free quantification (LFQ) values were compared using a Student’s T-test. For examining correct assignment of core RNP components (Figure 2b), RNPs were manually annotated using existing literature.^18,27–34^ The data were filtered to only include these components and mean LFQ values between replicates were collected. For each protein, the mean LFQ signal was normalized per protein by dividing by the maximum mean LFQ signal for that protein.

To assign distinct RNPs to each of the multi-RAP-MS targets, the LFQ values were analyzed by ANOVA. Significantly different proteins were selected using the Benjamini-Hochberg procedure (p < 0.05, FDR = 0.05). After removing non-significant proteins, protein were assigned to individual RNAs based on whether the mean LFQ was overrepresented for a given RNA target. The mean LFQ value across replicates was collected for each protein. These means were summed and the fraction of each mean was calculated (e.g. mean LFQ value of Snrnpg for U1 divided by the sum of the mean LFQ values of Snrnpg for all targets). If the protein was overrepresented in a given target, it was assigned to that RNA (e.g. for 8 targets, if the fraction of Snrnpg for U1 was greater than 1/8 of the summed mean signal for Snrnpg, Snrnpg would be assigned to U1). After the proteins were assigned to targets, any proteins not found in both replicates for that target were removed. The log10 ANOVA p-values and log10 mean LFQ values were used for Figures 2c and 3.

### RAP-MS 1.0

#### Probe design

U1 probes were identical to those used in RAP-MS 2.0 except that the 5’-CAAGTCA was replaced with a 5’-biotin. Probes were ordered from IDT with standard desalting.

#### RNA antisense purification followed by mass spectrometry (RAP-MS)

RAP-MS was performed as previously described.^17^ The only modification was that TCA precipitation, HiPPR purification, and HPLC desalting were replaced with the RAP-MS 2.0 Sp3-bead purification method.

### Covalent linkage affinity purification (CLAP)

#### Plasmid construction

Entry vectors for the targeted ORFs were ordered from DNASU and cloned into CLAP destination vectors (PYPP-CAG-Halo-V5-FLAG) using LR clonase. Correct inserts were validated through Sanger sequencing and validated plasmids were prepared with the ZymoPURE II Plasmid Midiprep Kit.

#### CLAP

CLAP was performed as previously described.^54,55^ Briefly, cells were lysed, and lysates were coupled to an agarose Halo resin. The resin was washed vigorously at 90°C under a variety of denaturing conditions. RNAs were freed from the resin using Proteinase K (NEB) and prepared into Illumina Sequencing Libraries. Libraries were sequenced on an Illumina NextSeq.

A custom mm10 genome containing repetitive elements and multicopy short RNAs was used for analysis as previously described.^83^ Reads were aligned using STAR and fold change over input was calculated using deepTools’s bamCompare tool.^84,85^

### iCLIP Reanalysis

#### Data analysis

FASTQ files for Alyref-FLAG, Chtop-FLAG, and Empty-FLAG iCLIP were pulled from GEO GSE113953.^53^ Reads were aligned using STAR to a previously described modified hg38 genome containing separate chromosomes for rRNAs and RMRP.^85,86^

### SPIDR Reanalysis

#### Data analysis

Differentially enriched second read truncations from published SPIDR data for Fus, Ewsr1, and Taf15 were plotted over the U1 snRNA gene body.

## BACK MATTER

## Acknowledgements

This study was supported by NIH grant P20GM125498 and UVM Cancer Center pilot grant (to DM).

## SUPPLEMENTAL FIGURES

**Supplemental Figure 1.**
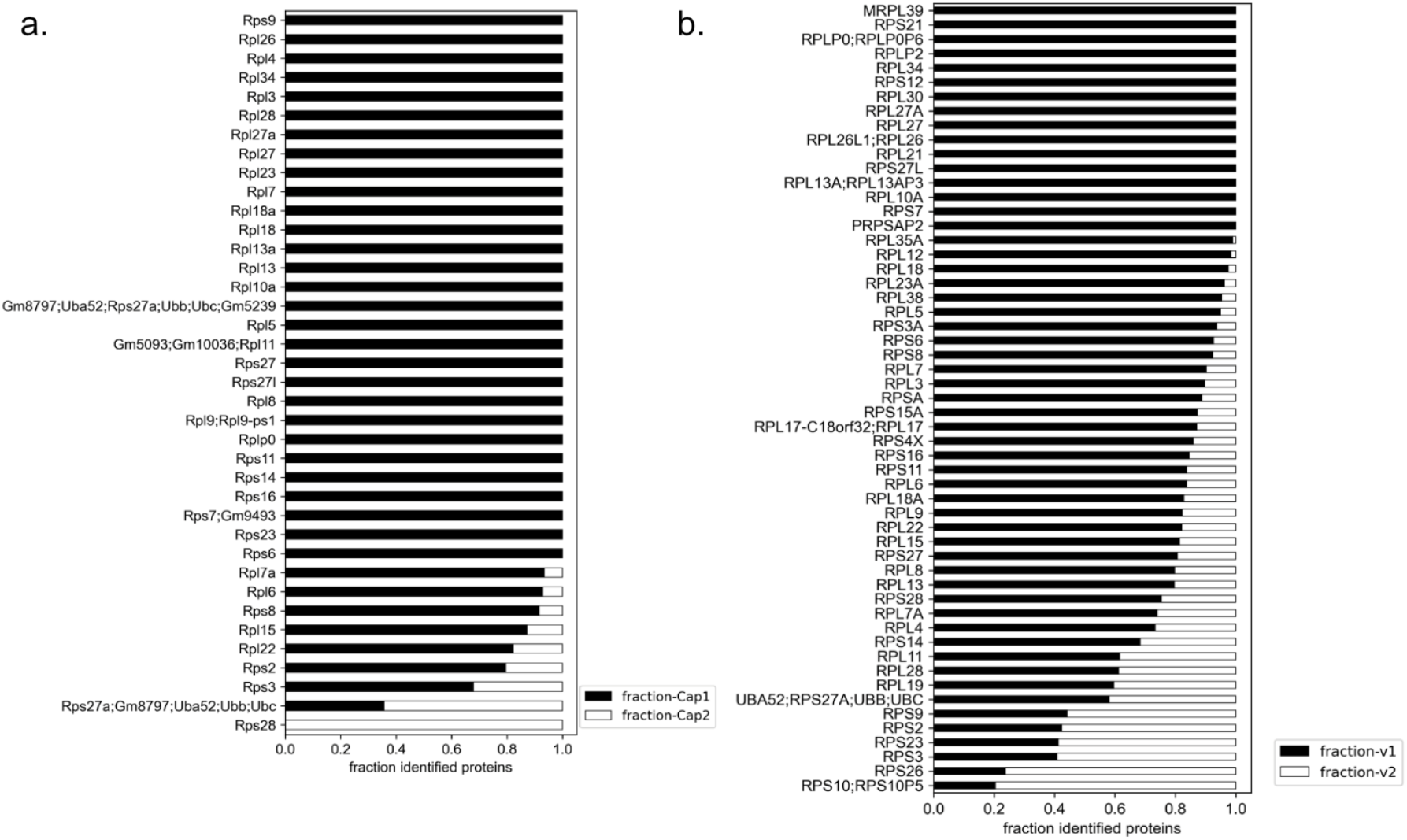
Ratios of ribosomal proteins in capture 1 and 2, and RAP-MS 1.0 and 2.0 comparison. a, Mean fraction LFQ for all ribosomal proteins identified in the U1 RAP-MS 2.0 experiment comparing captures 1 and 2. b, Mean fraction LFQ for all ribosomal proteins identified in the U1 RAP-MS experiment comparing our original protocol (RAP-MS 1.0) to the updated method presented in this paper (RAP-MS 2.0).

